# Cell type-specific monoclonal antibody cross-reactivity screening in non-human primates and development of comparative immunophenotyping panels for CyTOF

**DOI:** 10.1101/577759

**Authors:** Zachary B Bjornson-Hooper, Gabriela K Fragiadakis, Matthew H Spitzer, Deepthi Madhireddy, Kevin Hu, Kelly Lundsten, Garry P Nolan

## Abstract

Monoclonal antibodies are a critical tool for immunologists, with applications including immunomicroscopy and cytometry, and non-human primates are essential models for drug development. Very few antibodies are raised against non-human primate antigens; instead, researchers typically use anti-human antibodies that are found to be cross-reactive with non-human primates. The NIH maintains a valuable database of cross-reactivity, but its coverage is not complete, especially for less common species. Furthermore, the database only indicates the presence or absence of staining and does not indicate if a different cell population is stained in the NHP species than in humans. We screened 332 antibodies in five immune cell populations in blood from cynomologus macaques (*Macaca fascicularis*), rhesus macaques (*Macaca mulatta*), African green monkeys (*Chlorocebus aethiops*) and olive/yellow baboons (P*apio hamadryas anubis* × *Papio hamadryas cynocephalus*), thereby generating a comprehensive cross-reactivity catalog that includes cell type-specificity. We subsequently used this catalog to help create large CyTOF mass cytometry panels for three of those species and humans. The curated dataset containing the primary data for each antibody has been deposited as a browsable resource at https://immuneatlas.org and https://flowrepository.org/id/FR-FCM-Z2Z7.

## Introduction

Non-human primates (NHPs) are critical components of drug development because of their similarity to humans. Many key immunology assays, such as flow cytometry, Western blots, immunohistochemistry and immunofluorescence microscopy, make use of antibodies to demarcate specific cell types and quantify signaling moieties. Very few antibodies are raised against non-human primate antigens; instead, researchers typically use anti-human antibodies that are cross-reactive with the non-human primate species that they are studying. To help researchers find antibodies for NHP research, the National Institutes of Health supports a highly valuable database of the cross-reactivity of commercially available antibodies with 13 NHP species (http://www.nhpreagents.org). The database is derived from manufacturer and investigator reports, and typically provides a simple yes/no statement about whether a clone stains a species, with occasional comments about staining intensity or specificity. While an invaluable resource, the database is limited in its coverage. For example, prior to this study, only 28 CD markers had been evaluated in African green monkeys.

Additionally, with few exceptions, the database lacks information about the cell types bound by cross-reactive antibodies, and there are many known instances of antibody clones binding different cell types in different species. For example, granulocyte and monocyte marker expression is known to be substantially different in humans than in non-human primates. Anti-human CD33 clone AC104.3E3 was reported in the NIH database and manufacturer’s datasheet as cross-reactive with rhesus and cynomolgus macaques, but our lab and others determined that in those species, it prominently stains granulocytes (1, 2), while in humans it stains monocytes and classical dendritic cells. As another example, the Fcγ receptor CD16 is found on granulocytes in humans and sooty mangabeys, but not in macaques or baboons (3, 4), which will likely confound animal studies evaluating therapeutic antibodies, which may bind, transduce signals through and mediate internalization via this Fcγ receptor. Yet another example is CD56, which is expressed on monocytes in macaques (5), but is a canonical NK cell marker in humans. Thus, researchers must confirm that each clone they use is staining the cell population of interest through literature review or experimental verification.

Here we present an expansion of both the breadth and depth of primate cross-reactivity data. We screened 332 monoclonal antibodies in blood from two individuals of each of four NHP species: rhesus macaque (*Macaca mulatta*), cynomolgus macaque (*Macaca fascicularis*), African green monkey (*Chlorcebus aethiops*) and olive/ yellow baboon (*Papio hamadryas anubis* × *Papio hamadryas cyno-cephalus* hybrid); and found more than 120 clones that stained one or more populations in each species. Furthermore, we included counter-stain antibodies that allowed us to determine staining specificity in at least five major immune cell populations. Data from the NHPs were compared to the same screen ran in our laboratory on human blood. Finally, we used the results from this screen to create the first mass cytometry phenotyping panels for African green monkeys and rhesus and cynomolgus macaques. These panels allow for effective side-by-side comparisons between non-human primate and human blood. All data is publicly accessible at https://immuneatlas.org and https://flowrepository.org/id/FR-FCM-Z2Z7.

## Results

### Counterstain panel and staining condition development

In order to identify the characteristics of these antigens across species, we first designed a panel of counterstain antibodies that delineate major circulating immune cell types in all five species (Figure 1). This panel readily identified granulocytes, B cells, T cells, NK cells and monocytes/dendritic cells. In all species except for African green monkey, we could additionally separate the monocytes and dendritic cells based on CD11b expression. In African green monkeys, CD11b (ICRF44) was non-reactive, and we chose not to include a substitute marker to maintain technical consistency across all species.

**Figure 1.**
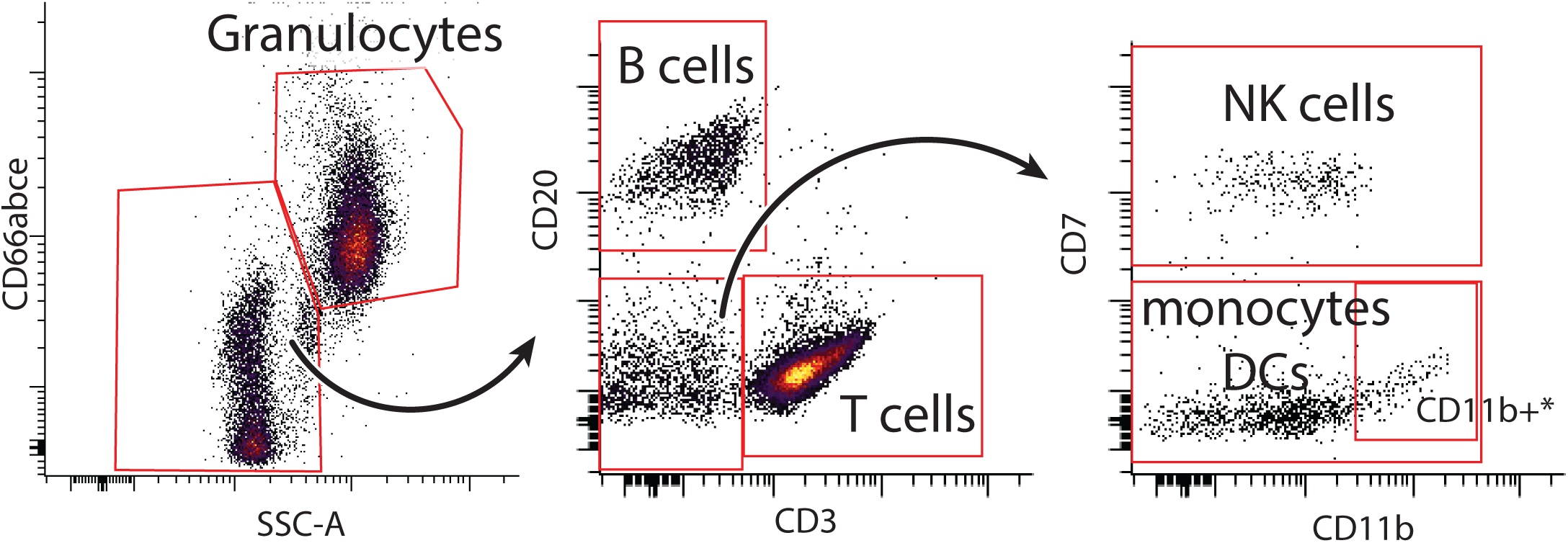
Five counterstains allow identification of at least five cell populations. Representative staining from a baboon shown. After gating by time to exclude artifacts, granulocytes were identified as CD66abce+/SSC-A+. Non-granulocytes were divided into B cells (CD20+/CD3-), T cells (CD3+/CD20-), NK cells (CD7+/CD3-/CD20-) and monocytes/dendritic cells (CD7-/CD3-/CD20-). *In species other than the African green monkey, dendritic cells could be separated on the basis of CD11b staining.

Additionally, we strategically selected the fluorophores to keep the PE channel, which was used for the screened antibody, free of bleed/compensation to avoid technical artifacts.

Fixation and red blood cell lysis conditions were selected after testing a large number of protocols on the basis of preserving antigen staining and effectively lysing non-human primate blood (data not shown). Fixation with approximately 0.3% PFA was low enough to avoid obvious loss of staining and sufficient to minimize morphological changes of cells as observed as forward and side scatter signals changing over the course of acquisition on the cytometer. Enzymatic lysis with VersaLyse was highly effective for both human and non-human primate blood, whereas other methods such as hypotonic lysis were inadequate or inconsistent for non-human primate blood.

### Antibody reactivity screen

We established universal criteria for defining positive expression to determine antigen expression in an unbiased manner. An antibody clone was classified as reactive with a population if more than 10 percent of cells had a PE signal intensity greater than the 95^th^ percentile of the intensity of the corresponding isotype control for the same species and population (Table I). This threshold was found to accurately reflect the results of manual classification: We verified 500 of the 14,940 clone × species × population results, taking into consideration reported staining patterns (references included (2, 5, 7–13)) and the visual degree of separation from the isotype control, and calculated a false-positive rate of 7.4% and a false-negative rate of 1.6%, for an initial accuracy of 91% in our data. Then, we manually verified and corrected as necessary all discordant replicates and all clones that were classified as reactive in a non-human primate species but not in a human; thus, our final estimated accuracy exceeds 91%. All data from this project are publicly available, such that researchers can independently validate the expression of antigens of interest by accessing the primary data. In total, we identified 260, 153, 129, 161 and 147 clones that are reactive with one or more populations in human, cynomolgus macaque, rhesus macaque, African green monkey and baboon, respectively.

**Table I:**
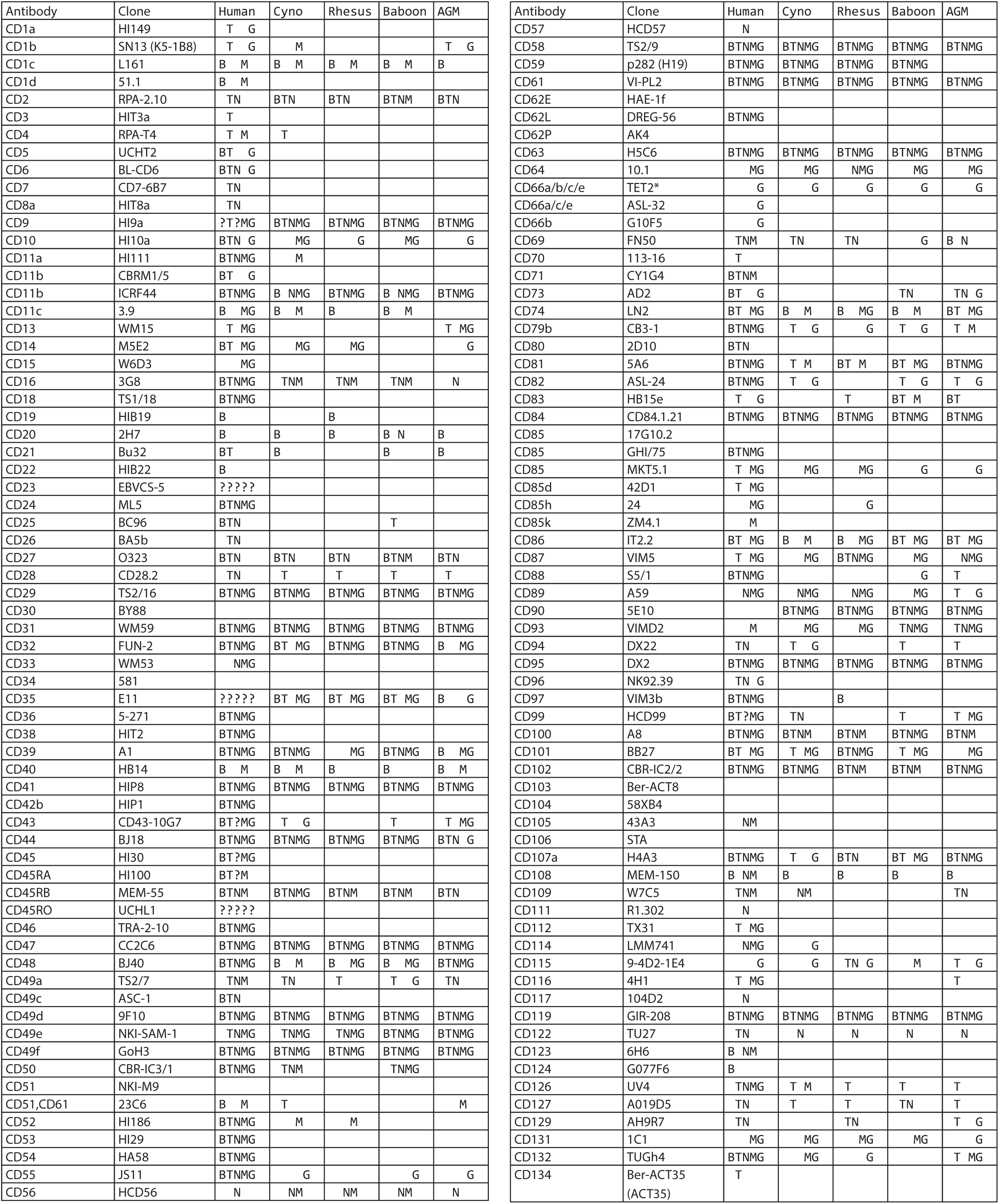
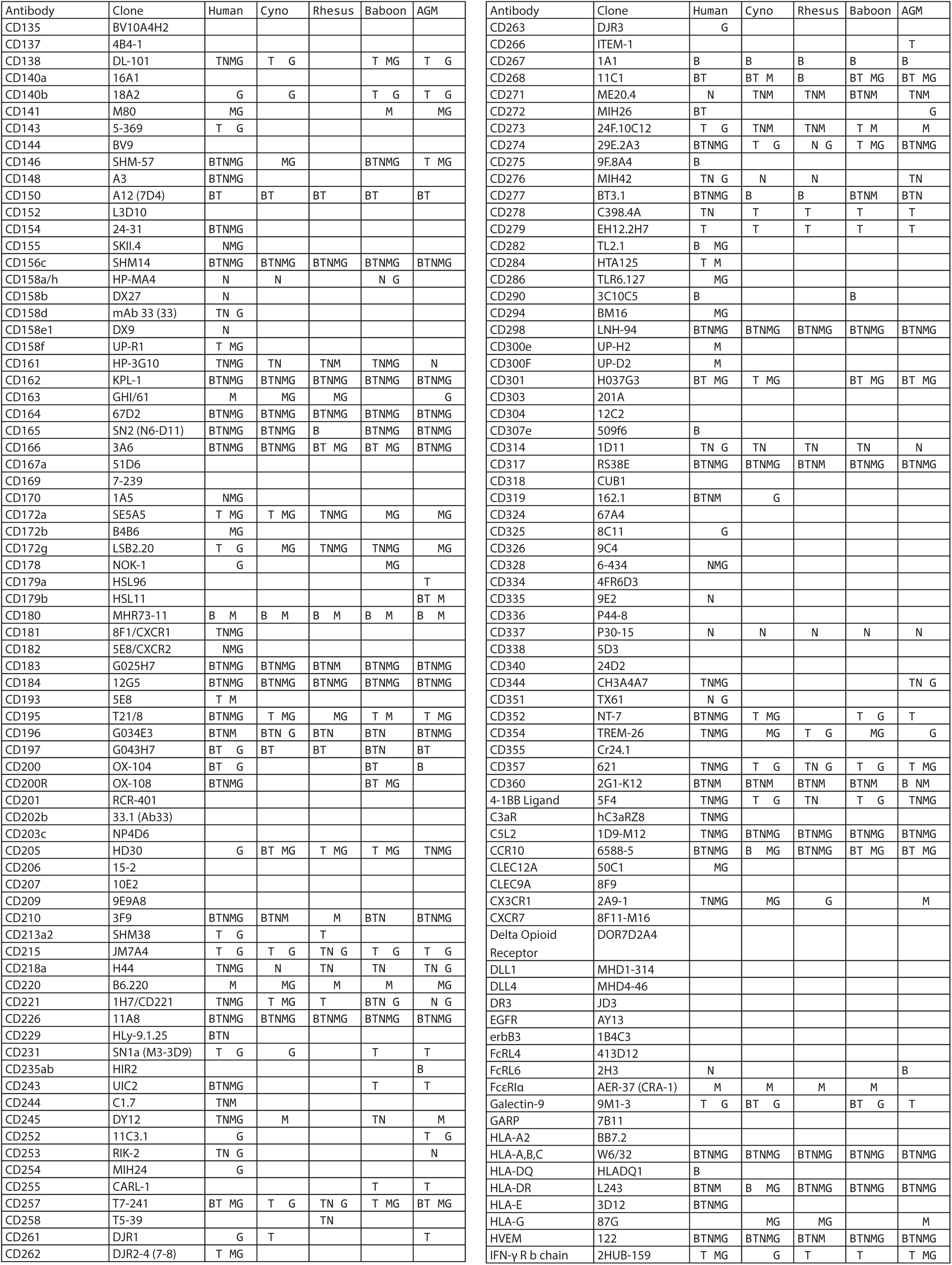
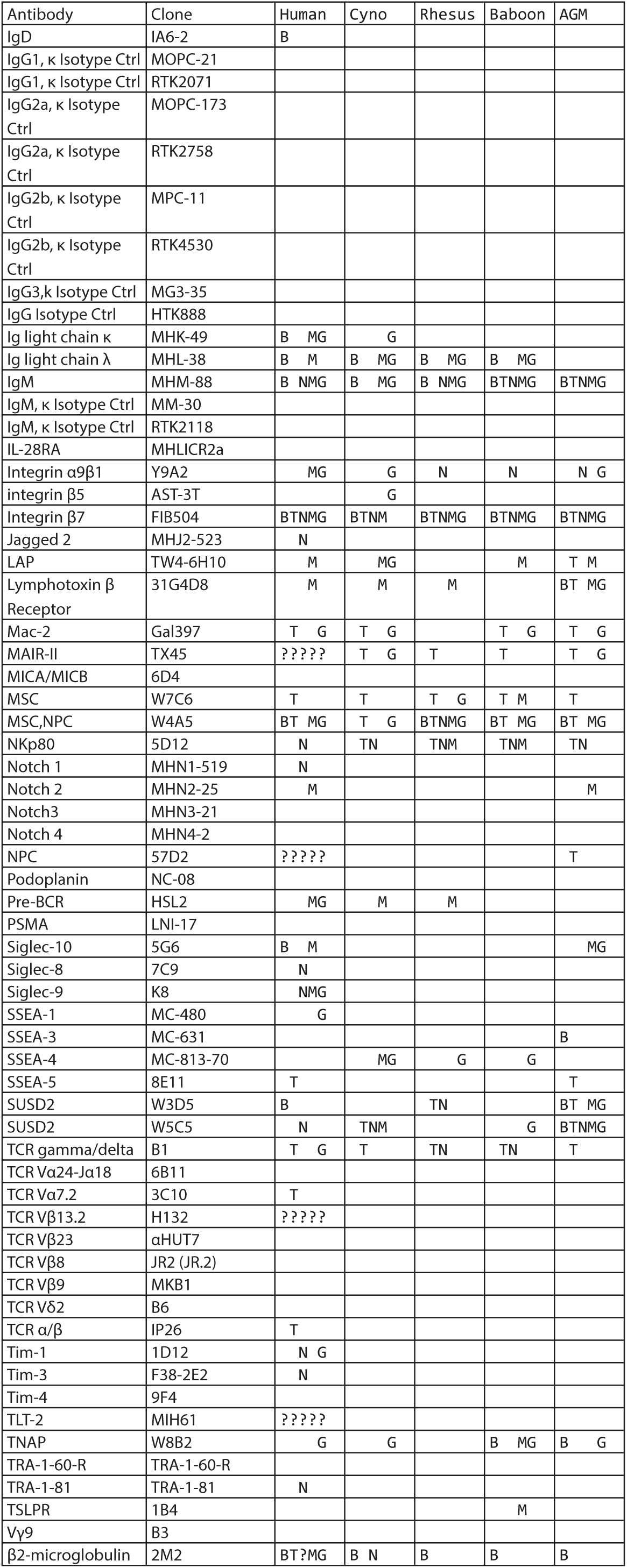
Summary of cell type-specific cross-reactivity. Clones were considered reactive if at least 10 percent of cells had a signal greater than the signal of the 95th percentile of corresponding isotype control or if they were manually classified as positive. Presence of letter in column indicates reactive staining detected in that cell population, absence of letter indicates clone is unreactive in that cell population. B: B cells, T: T cells, N: CD7+ NK cells, M: CD7-monocytes and dendritic cells, G: granulocytes, ?: inconclusive, insufficient number of cells acquired for a given population to make clear assessment. Two of each non-human primate species and one human were assayed.

Researchers interested in evaluating the antibodies in this study are directed to examine the primary data at https://immuneat-las.org or https://flowrepository.org/id/FR-FCM-Z2Z7, where one can compare the relative staining intensity and distribution of staining between species. Such review is critical when evaluating antibodies that stain rare populations—especially populations that comprise less than 10 percent of one of the populations that we delineated, or that show dim expression that might be excluded by the threshold for expression and staining that we applied. Additionally, we encourage diligence when interpreting markers such as CD41 and CD51/ CD61, which are listed as reactive with all cell types, but in actuality are probably staining platelet fragments stuck to other cells, based on the known distribution of those markers in humans (14). It is also important to keep in mind that actual antibody-antigen specificity may vary by species due to differences in gene sequence, protein structure and post-translational modifications.

### Notable examples of differences in expression patterns

As noted above, in macaques CD33 is found on granulocytes and CD16 is restricted to monocytes and dendritic cells. In this study, we found that African green monkeys share this same staining pattern, albeit with weaker CD33 staining. Another notable difference (out of numerous idiosyncratic expression patterns observed) is that CD172g (signal regulatory protein (SIRP) γ, also known as SIRPβ2) is expressed on CD11b+ monocytes and granulocytes, but not on T cells, in all of the NHP species examined. In contrast, in humans this marker is expressed on T cells, some B cells and to some degree in granulocytes (Figure 2). Because it lacks a cytoplasmic signaling domain, CD172g is postulated to signal unidirectionally by activating the CD47-expressing cell and inducing T cell migration and proliferation in humans, mice and rats (15–17). Thus, our finding suggests a major difference in regulation of immune cell migration and adaptive response, which warrants further study.

**Figure 2.**
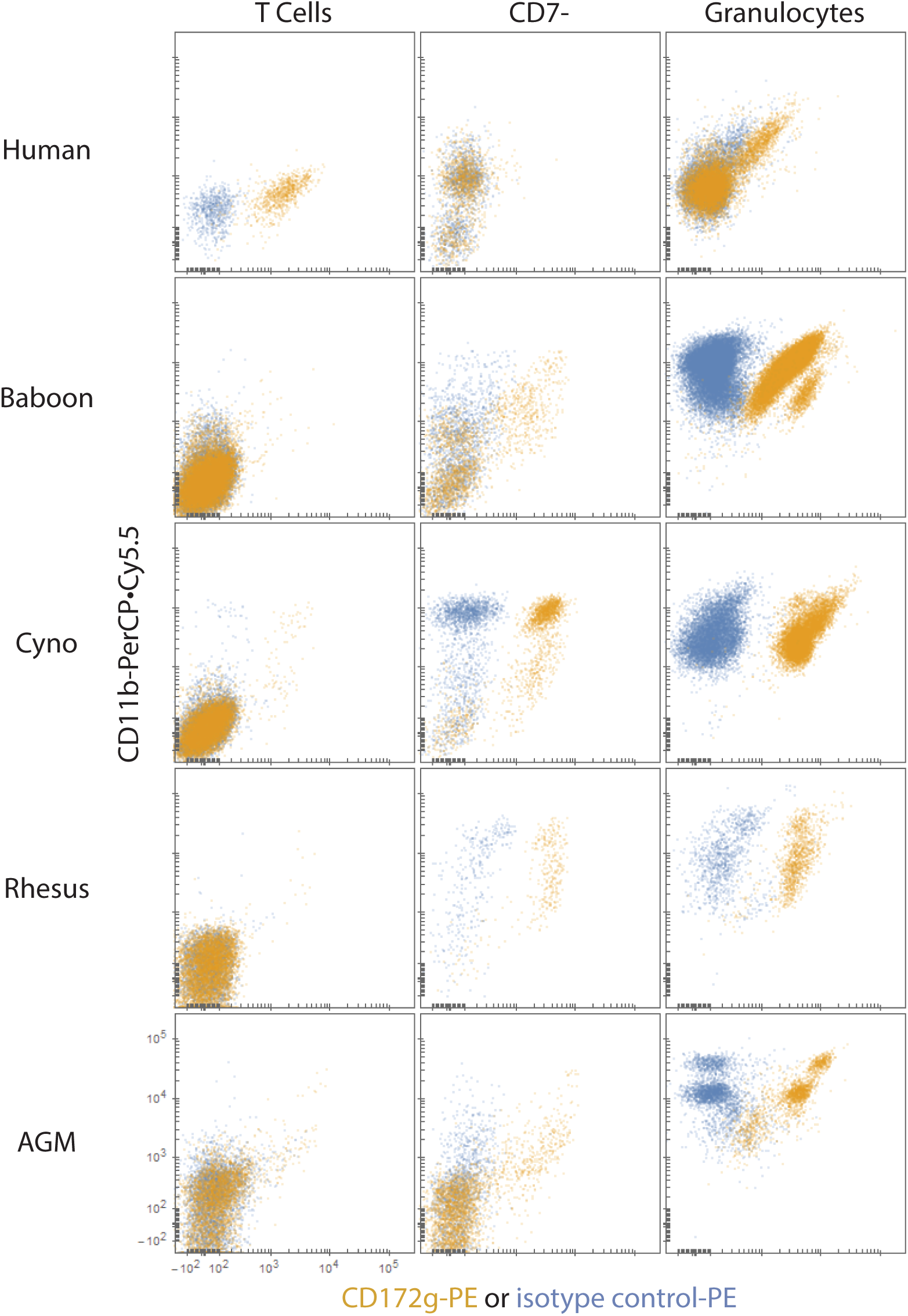
CD172g is expressed on monocytes and all granulocytes, but not on T cells, in examined NHP species. By comparison, CD172g is expressed on all T cells, a subset of granulocytes and a subset of B cells (data not shown) in humans. As discussed in the text, CD11b did not uniformly stain AGMs; thus, some of the CD11b-negative cells are monocytes. Blue: isotype control, orange: CD172g.

Another notable example of the differences found is the presence of CD2 staining not only on T cells, but also on B cells in rhesus macaques, cynomolgus macaques, African green monkeys and to a slight extent baboons (Figure 3). CD2 is involved in adhesion, co-stimulation, antigen recognition and potentially differentiation (18, 19). In mice, virtually all circulating and bone marrow B cells express CD2 (18). Kingma et al. previously reported that only a small subset (5.74 +/-2.46%) of normal human peripheral blood B cells express CD2, although approximately 25% of surveyed B cell neoplasms expressed CD2 (20). We observed no CD2+ B cells in human. B cell CD2 expression thus seems to have been lost evolutionarily, with the most abundant expression in the oldest species (mouse), moderate expression in macaques and African greens, less expression in baboons and essentially no expression in the youngest species (human). Interestingly, the ligand is not conserved between species: CD2 exclusively binds CD48 in mice and rats, while in humans it strongly binds CD58 and only weakly binds CD48, which is also co-expressed with CD58 (19, 21).

**Figure 3.**
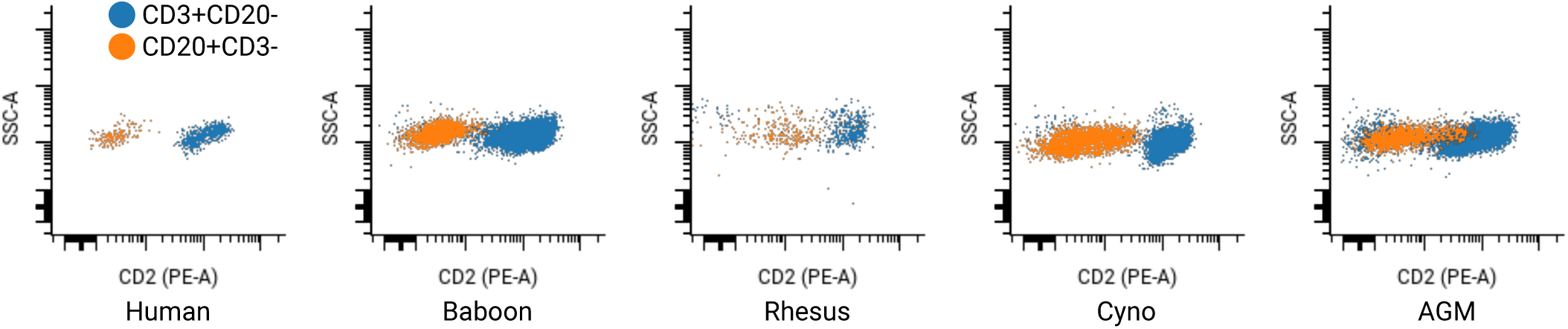
CD2 is expressed on B cells in NHPs. Blue: CD3+ T cells where expression is expected in humans; orange: CD20+ B cells.

### CyTOF panel design

We used the results of the antibody screen, along with the results of several targeted follow-up experiments, to craft a set of parallel CyTOF mass cytometry panels for rhesus and cynomolgus macaque, African green monkey and human (Table II).

**Table II:** CyTOF mass cytometry panels for orthologous phenotyping of humans and three species of NHPs. Canonical cell targets (rows) are listed along with mass channel, and antibody clones used for staining humans, macaques, and African green. Clones are listed in parentheses. * indicates weak staining.

| Canonical Target | Channel | Human | Macaque | African green |
| --- | --- | --- | --- | --- |
| Lymph/mono | 115 | CD45 (HI30) | CD45 (DO58-1283) | CD45 (DO58-1283) |
| Erythrocytes | 113 | CD235a (HIR2) | CD233 (BRIC6) |  |
| Platelets | 140 | CD61 (VI-PL2) | CD61 (VI-PL2) | CD61 (VI-PL2) |
| Granulocytes | 158 | CD66abce (TET2) | CD66abce (TET2) | CD66abce (TET2) |
| Granulocytes | 142 |  | CD33 (AC104.3E3) | CD33 (AC104.3E3)* |
| B cells | 173 | CD20 (2H7) | CD20 (2H7) | CD20 (2H7) |
| B cells, class-switched | 174 | IgM (G20-127) | IgM (G20-127) | IgM (G20-127) |
| APCs (B, DC) | 176 | HLA-DR (Immu357) | HLA-DR (Immu357) | HLA-DR (Immu357) |
| DCs, B cells | 162 | CD1c (AD5-8E7) | CD1c (AD5-8E7) | CD1c (AD5-8E7) |
| cDCs | 164 | BDCA3 (1A4) | BDCA3 (1A4) | BDCA3 (1A4) |
| pDCs | 148 | CD123 (7G3) | CD123 (7G3) | CD123 (7G3) |
| T cells | 157 | CD3 (SP34.2) | CD3 (SP34.2) | CD3 (SP34.2) |
| CD4+ T cells | 156 | CD4 (OKT4) | CD4 (OKT4) | CD4 (L200) |
| CD8+ T cells | 155 | CD8a (RPA-T8) | CD8a (RPA-T8) | CD8a (RPA-T8) |
| Naïve T cells | 166 | CD45RA (HI100) | CD45RA (HI100) |  |
| Myeloid cells | 153 | CD11b (ICRF44) | CD11b (ICRF44) | CD11b (ICRF44) |
| Myeloid, T subset | 143 | CD11c (Bu15) | CD11c (3.9) | CD11c (3.9) |
| Monocytes | 151 | CD14 (M5E2) | CD14 (M5E2) | CD14 (M5E2) |
| NK, mono, gran | 159 | CD16 (3G8) | CD16 (3G8) | CD16 (3G8) |
| Myeloid | 142 | CD33 (AC104.3E3) |  |  |
| Monocytes | 175 |  | CD56 (NCAM16.2) |  |
| NK cells | 175 | CD56 (NCAM16.2) |  |  |
| NK cells | 141 | CD7 (M-T701) | CD7 (M-T701) | CD7 (M-T701) |
| NK cells | 168 | CD161 (HP-G310) | CD161 (HP-G310) | CD161 (HP-G310) |

Wherever possible, we used the same antibody clones for all species. In the case of erythrocytes, we used the common marker CD235a for humans and CD233 for NHPs, instead of using the less common CD233 for all species (no anti-CD235 clone reactive with NHPs could be found). In the case of CD11c, clone 3.9 was only used in non-human primates because Bu15 was non-reactive, but 3.9 is reported to preferentially bind activated CD11c (22) and requires the addition of magnesium as a cofactor during staining (22, 23). We primarily use CD16 in lieu of CD11c during gating analysis for consistency. In the case of naïve and memory T cells, we observed very broad distribution of CD45RA staining, as previously reported (24, 25), with two clones (5H9 and HI100), which was difficult to gate (see online dataset). Staining of this marker was superior with the Versa-Lyse-based method than with the Triton-X100-based method (see methods) (data not shown).

The experiments described here are especially valuable for the advancement of African green monkey (AGM) immunology because there is a dearth of literature discussing immunophenotyping and only 28 reactive antibodies listed in the NIH database. Nonetheless, this species is important in drug development—especially SIV research—and is the subject of an international effort to make it the most comprehensively characterized NHP by phenotype and genomics (26). Notably, the usage of AGMs is increasing due to a shortage of rhesus macaques for research (27). It was generally possible to use the same clones for African green monkeys as we were for macaques, with several exceptions (Table II). No reactive clones were found for CD11c; instead, we again rely on CD16 for gating monocyte subpopulations and CD123, CD1c and BDCA3 for gating dendritic cell subpopulations.

These CyTOF panels were exhaustively titrated to determine optimal staining concentrations for each antibody, then revalidated in at least two donors from each species. We subsequently contracted BioLyph LLC to lyophilize and package several thousand single-use pellets (“LyoSpheres”) of each of these panels for use in later studies. These pellets solve stability issues such as evaporation and precipitation that inhibit long-term studies and eliminate the technical variability and errors that arise from pipetting. As such, these panels allow for parallel evaluation of orthologous populations between species (Figure 4).

**Figure 4:**
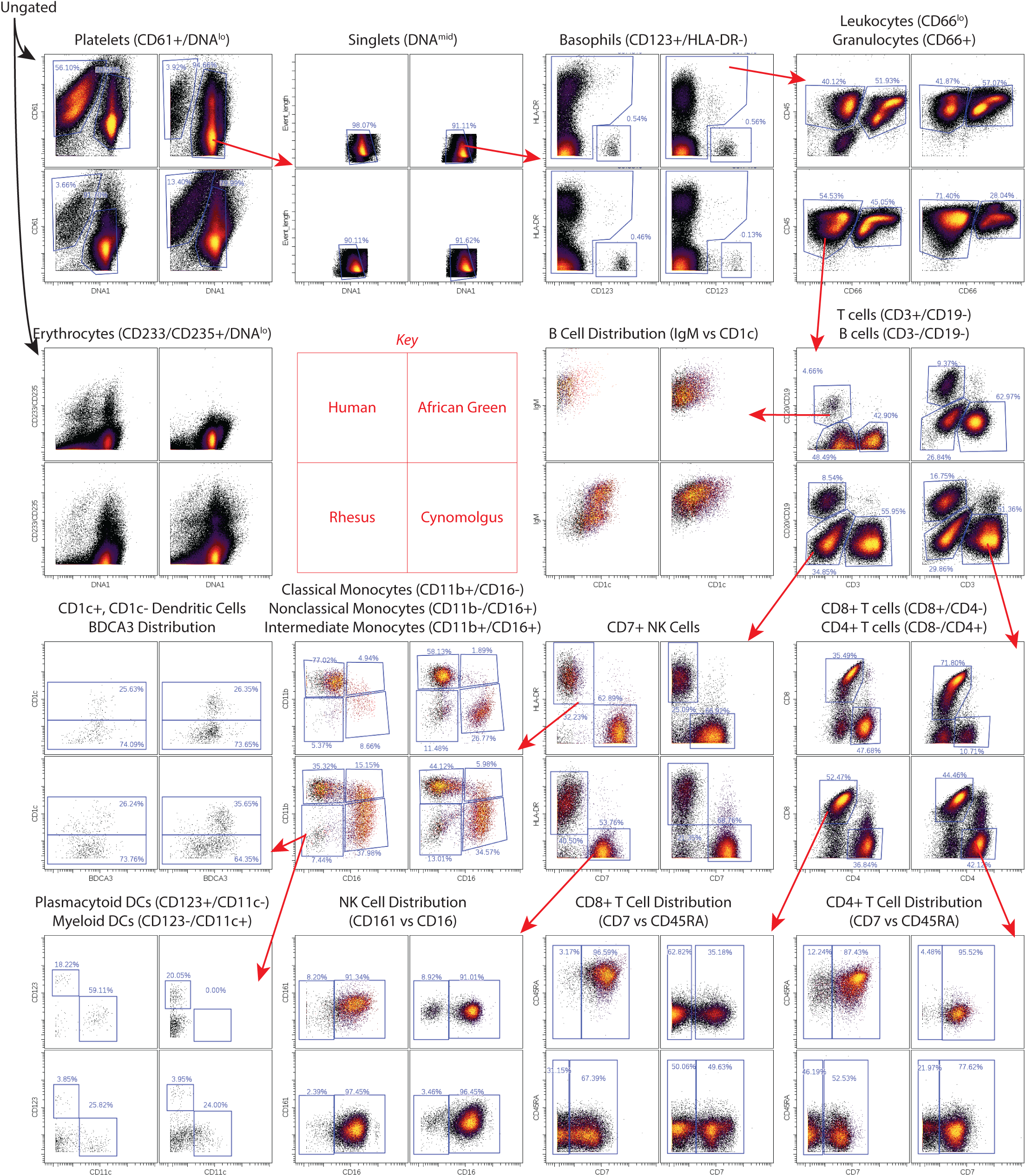
Parallel gating of blood from four species. CyTOF mass cytometry panels were developed to delineate at least 18 orthologous populations between species.

## Discussion

We have substantially expanded the breadth and depth of available antibody cross-reactivity data between primate species, and have deposited the dataset containing the primary flow cytometry data online so that any user can easily compare staining patterns in these species, and even look at subpopulations of cells to check specificity—information not typically reported in the NIH database.

As a result of evaluating specific cell types, we have identified numerous antibodies that stain different populations in non-human primates than in humans. In light of this, researchers must take adequate steps to ensure they are evaluating the intended populations when using these reagents, and when evaluating immunotherapeutics targeting these proteins.

While we screened 332 different anti-human antibody clones, not all of the antigens targeted by those antibodies are expected to be present in the resting, peripheral blood cells that we tested. Indeed, only 78.3% (260/332) of the antibodies positively stained human blood. Thus, our screen did not evaluate markers that are found only in cells from bone marrow and other tissues, in progenitors or in activated populations. However, blood is the most easily accessed and likely the most common biological sample type used in research environments, and we thus expect this knowledge about blood to be of the greatest use to most researchers. We hope that the framework we have developed will be expanded upon with additional clones and/or additional species in future studies.

Finally, we present the first CyTOF mass cytometry panels for coordinated human and non-human primate research. We have continued to use these panels for large-scale comparisons of primate immune systems (in review).

## Materials and Methods

### Blood

Macaque blood was obtained from Valley Biosystems, Inc. (location withheld). African green monkey blood was obtained from Bioreclamation, LLC (Westbury, NY) and Worldwide Primates, Inc. (location withheld). Baboon blood was obtained from the Southwest National Primate Research Center, which is funded by the National Center for Research Resources (p51 RR013986) and supported by the Office of Research Infrastructure Programs/OD P51 OD011133. All animal blood was collected under an approved animal care and use protocol. Human blood was obtained from a healthy adult donor under an approved IRB protocol.

### Antibody screen

Blood was collected in sodium heparin and processed within 24 hours of collection. Eight to 10 ml of blood was gently fixed and lysed by incubating with 792 µl of 16% paraformaldehyde (Electron Microscopy Sciences, Hatfield, PA; final concentration approx. 0.3%) and 29.6 ml of VersaLyse (Beckman Coulter, Brea, CA) for 10 minutes at room temperature, then washed once with 0.01% BSA in PBS (“staining buffer”). Cells were resuspended in 6.5 ml of staining buffer, then incubated with 0.5 ml of human TruStain FcX Fc receptor blocking solution (Biolegend, San Diego, CA) for 10 minutes.

Five counterstain antibodies were then added: 500 µl CD3 (SP34.2)-Brilliant Violet 421 (BD Biosciences, San Jose, CA), 500 µl CD20 (2H7)-Brilliant Violet 605 (Biolegend), 1000 µl CD66 (TET2)-APC-Vio770 (Miltenyi, San Diego, CA), 500 µl CD11b (ICRF44)-PerCP/ Cy5.5 (Biolegend) and 500 µl CD7 (M-T701)-APC (BD Biosciences), for a final volume of 10 ml.

Subsequent steps were performed on an automation platform including an Agilent Bravo 96-channel pipetting robot, centrifuges, BioTek ELx405 96-channel aspirator/dispenser and Thermo Scientific MultiDrop dispenser. Twenty microliters of cells in the antibody cocktail were dispensed into every well of a 384-well plate using the MultiDrop. LegendScreen Human PE antibody screen plates (Biolegend) were rehydrated with the manufacturer-recommended 25 µl of water. The screen consists of four 96-well plates; 5 µl from each well were transferred to the single 384-well plate quadrant-wise. Staining reactions were incubated in the dark at room temperature for 30 minutes with 2.0 mm-radius orbital shaking, then washed three times with staining buffer and acquired on a BD LSR II with 405, 488 and 633 nm lasers using an HTS autosampler.

Files were gated and populations exported using a web-based, high-throughput cytometry analysis platform developed in our laboratory. Every well was manually gated by time to exclude anomalies caused by air bubbles or debris: temporal regions where the signal in the PE channel over time were inconsistent were excluded. Cell populations were then gated as described in the results section. Population gates were tailored to each species; tailoring to individual donors within a species was not necessary. The remaining analysis (statistics calculations, discordant replicate resolution, verification and reporting) was performed using Mathematica (Wolfram Research, Champaign, IL).

### CyTOF mass cytometry

Purified antibodies (BioLegend, BD, Beckman Coulter) were conjugated to stable metal isotopes using commercially available polymeric chelators (“X8,” Fluidigim, Markham, Ontario, Canada). Whole blood was lysed and prepared using the method of Chow and Hedley (6), which is better suited to high-throughput, plate-based assays than the Versalyse method described earlier. Briefly, we mixed 330 µl of blood with 55 µl of 16% PFA, incubated for 10 minutes, then added 1,615 µl of 0.127% Triton-X100 (Fisher Scientific, Waltham, MA) in PBS (final concentration of 0.1% Triton) and incubated for 30 minutes. We then washed, stained with the metal-tagged antibodies for 30 minutes and washed again. Cells were subjected to a second fixation with 1.5% paraformaldehyde and DNA staining with iridium intercalator (Fluidigm), then acquired on a CyTOF 2 mass cytometer (Fluidigm).

## Disclaimer and Acknowledgements

The research discussed in this article was supported in part by the U.S. Food and Drug Administration (Contract No. HHSF223201210194C). Additional support was provided by NIH awards 5R01CA18496804, 5R25CA18099304, 1R01GM10983604, 5UH2AR06767603, 1R01NS08953304 and R01HL120724, and FDA contract HHSF223201610018C. Z.B.B.H. was supported in part by NIH grant T32GM007276. G.K.F. was supported in part by a Stanford Bio-X graduate research fellowship and NIH grant T32GM007276. M.H.S. was supported in part by NIH grant DP5OD023056. This article reflects the views of the authors and should not be construed to represent the U.S. Food and Drug Administration or NIH’s views or policies.

Thanks goes to BioLegend Inc. for providing the screening plates, and to the NIH Nonhuman Primate Reagent Resource (R24 RR016001 and NIAID contract HHSN 2722000900037C).

### Author contributions

Z.B.B.H. generated the data, performed the analysis and wrote the manuscript. G.K.F. and M.H.S. contributed to panel design and edited the manuscript. K.H. contributed to data generation. D.M. contributed to data generation and reagent optimization. K.L. edited the manuscript and advised analysis. G.P.N. advised the study and edited the manuscript.

